# A Look-ahead Monte Carlo Simulation Method for Improving Parental Selection in Trait Introgression

**DOI:** 10.1101/2020.09.01.278242

**Authors:** Saba Moeinizade, Ye Han, Hieu Pham, Guiping Hu, Lizhi Wang

## Abstract

Multiple trait introgression is the process by which multiple desirable traits are converted from a donor to a recipient cultivar through backcrossing and selfing. The goal of this procedure is to recover all the attributes of the recipient cultivar, with the addition of the specified desirable traits. A crucial step in this process is the selection of parents to form new crosses. In this study, we propose a new selection approach that estimates the genetic distribution of the progeny of backcrosses after multiple generations using information of recombination events. To demonstrate the effectiveness of the proposed method, a case study has been conducted using maize data where our method is compared with state-of-the-art approaches. Simulation results suggest that the proposed method, look-ahead Monte Carlo, achieves higher probability of success than existing approaches.

## 1 Introduction

From a commercial breeding perspective, trait introgression (TI) is a necessary process to produce the elite cultivar with the most desirable traits (1). Oftentimes introgressing favorable quantitative trait locus (marker) alleles from one population possessing desirable traits may benefit another crop population (2). As an illustration, imagine two maize populations: one population (recipient) characterized by high yielding potential and low resistance to drought stress, whereas the other population (donor) characterized by low yield potential and high resistance to drought stress. In this scenario, one would hope to recover all the attributes of the recipient while obtaining the drought resistant alleles of the donor by some mechanized breeding process to create a new elite cultivar. However, due to uncertainty in the breeding process, the new resulting cultivar will need to go through a rigorous selection process to determine if both desired traits have been inherited, thus, adding additional time to the process.

Although, in principle, the intent of trait introgression is forthright, in practice, there exists many complications due to the stochastic nature and size of a commercial breeding program. Because of this uncertainty, multiple breeding generations may be required until the superior, desired cultivar is achieved (3). An additional challenge of the TI process is deciding the parental crosses to perform out of a sizable, complex gene pool. At each generation cycle, plant breeders are faced with the difficult decision of identifying crosses to perform to produce the next generation of, hopefully, superior cultivar out of this large gene pool. In the prefect scenario, plant breeders would be able to cross every possible combination of parents until the desired cultivar is achieved. However, in reality, due to the limited amount of available resources (time, money, land, technology, etc.), breeders may only consider a small fraction of an existing gene pool, possibly leading to sub-optimal decision making (4; 5; 6). If only a single allele is needed, molecular assisted introgression can achieve this task. Whereas in the case of multiple desirable alleles, more complicated gene strategies will be necessary (7; 8; 9).

Recent advances in simulation and optimization techniques have been applied to help aid in the breeding process. Computer simulation approaches help identify optimal breeding strategies by adopting assumptions of the breeding system and running multiple scenarios, whereas, optimization approaches aim to produce the best framework to maximize the probability of achieving the desired cultivar while minimizing input resources. It should be noted that the combination of analytical techniques and plant breeding has mainly been applied to genomic selection and not trait introgression (10; 11; 12; 13; 14; 15).

Although there does not currently exist much literature to integrate operations research techniques and trait introgression, there are still a few impactful studies. Cameron et al. utilized an operations research framework with a stochastic optimization model to identify the best breeding strategies for a given population under resource constraints (16). This work illustrates the potential optimization modeling can have on resource allocation in plant breeding. Probabilistic simulation techniques have also been performed by Sun et al. to assess *in silico* various crossing schemes and breeding approaches (17). Moreover, Han et al. has framed trait introgression as an algorithmic process and introduced a novel selection metric, predicted cross value (PCV), which predicts specific combining ability by estimating the probability that a pair of parents will produce a perfect gamete with all desirable alleles (18).

Due to the importance of optimizing the breeding pipeline and the need to consider resource limitations for large scale breeding programs, this paper aims to design a platform that integrates operations research methods to trait introgression. Specifically, the authors develop a novel Monte Carlo simulation approach for the TI pipeline to consider the parental selection aspect under different scenarios of resources present within a commercial scale TI program. The originality in the proposed method, look-ahead Monte Carlo (LMC), is to look-ahead and estimate the performance of progeny in the target generation and then optimize the selection decisions based on the estimated performance. In the remaining of this paper, we first introduce two existing selection approaches and then compare the proposed method against the existing approaches in a case study using realistic maize data.

## 2 Methods

Backcrossing is a well-known breeding approach that can be employed to introduce a specific trait, such as disease resistance, from one individual, often an unimproved one, to another individual that is typically an elite breeding individual. The donor parent (DR) provides the desired trait and may not perform as well as an elite variety in other areas. The elite line, called the recurrent parent (RP), usually performs well in the background. The goal of backcrossing is to recover all the attributes of the recipient cultivar, with the addition of the specified desirable traits. First an initial cross is made between the donor and recurrent parent to produce F1 progeny. Since, the donor and recurrent parents are both homozygous, this step is deterministic which means the F1 progeny has 50% of the genetic material from each parent (Fig. 1). Next, the F1 individual is crossed to the recurrent parent to develop a backcross one (BC1) population. In Figure 1, we see *n* individuals in the backcross one population denoted with BC1_1_, BC1_2_,…, and BC1_*n*_. Individuals from the BC1 population will be selected based on a predefined metric and then again crossed to the recurrent parent.

**Figure 1.**
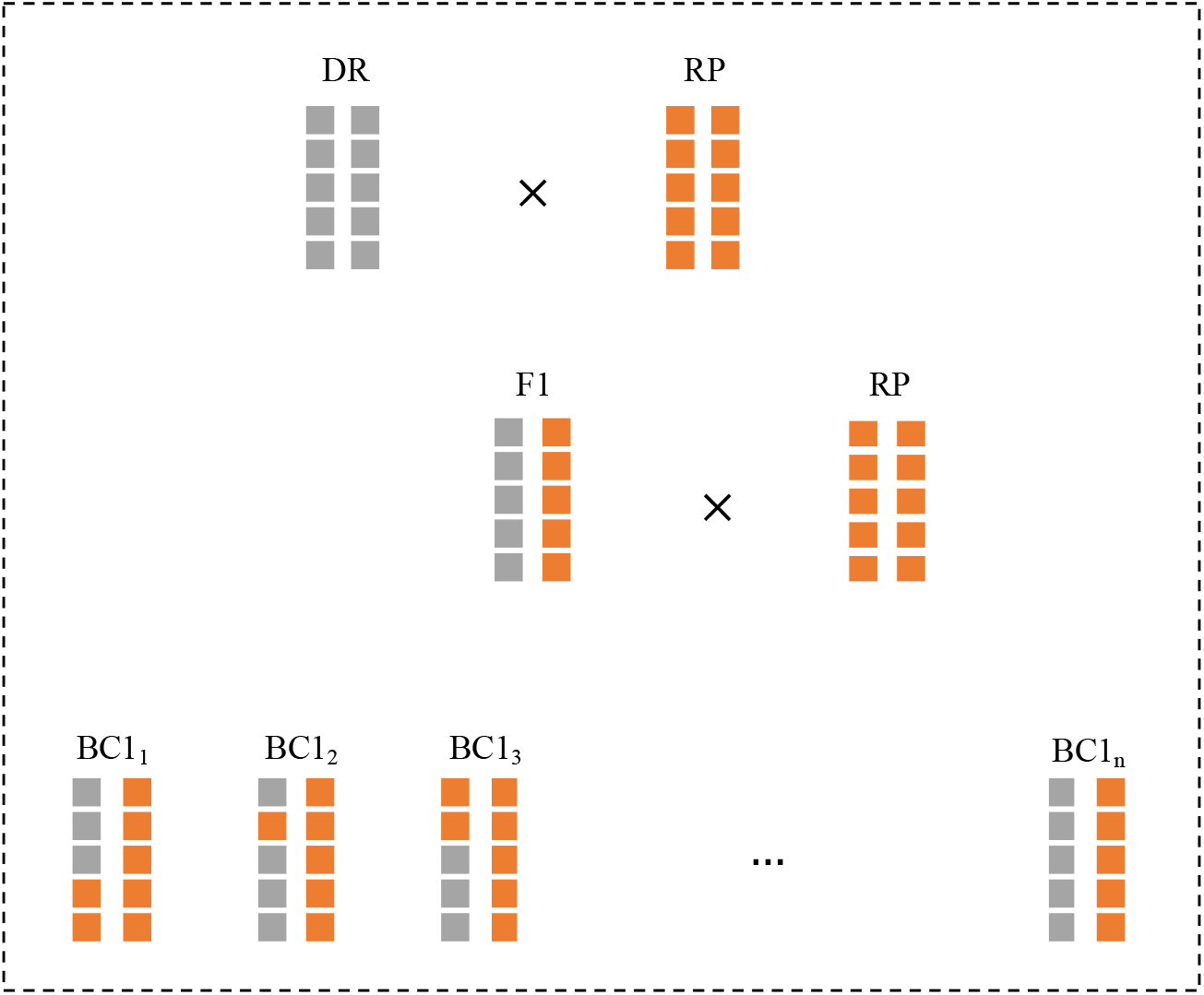
The donor parent is crossed with the recurrent parent to produce F1. F1 progeny have 50% of their genetic material from each parent. Then, F1 is backcrossed with the recurrent parent to develop the BC1 population. Here, BC1_*i*_ represents the *i*^th^ individual in the backcross one population.

In successive generations, progeny are first selected for the trait of interest and then backcrossed to the recurrent parent. This cycle continues until a new line that is highly close to the recurrent parent, but with the desired alleles or traits from the donor parent, is created. However, a crucial step in this process is the selection of parents to form new crosses. In the remainder of this section, we first describe two existing selection methods and then propose a new, novel selection approach that predicts the genetic distribution of the progeny of backcrosses after multiple generations using information of recombination events.

### 2.1 Background selection

The background selection approach first selects the individuals with desired marker genotypes and then among these positive individuals selects for the desired background genotype (19; 20; 21). Background selection has been shown to be efficient by previous theoretical work (19; 20; 22; 23) and experimental work (24).

The breeding value of a background genotype can be estimated using genomic estimated breeding value (GEBV) (25; 26). To represent the genotype of an individual plant, we use an *L × m* binary matrix, say *G* ∈ **B**^*L×m*^, where *G_l,m_* = 1 indicates whether the *l*^th^ allele from chromosome *m* is desirable or not (*G_l,m_* = 0). For each individual plant represented with a binary matrix, *L* is the total number of markers in the genome and each row is a locus in the genome. The number of columns in the binary matrix represents the ploidy of the plant. We use diploid species in this paper (*m* = 2). Additionally, we assume **D** = {*d*_1_, *d*_2_,…, *d_z_*} is the location of the positive markers from the donor and there are total *Z* markers that should be introgressed. If we assume uniform weight for all desirable alleles, then the background GEBV of an individual is equivalent to the number of desirable alleles in its background:

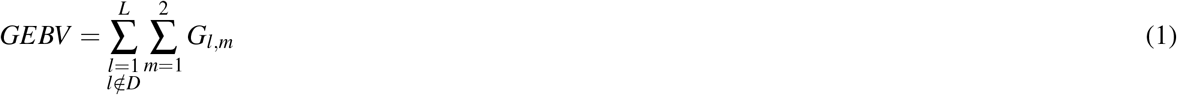

According to this approach, the positive individuals with highest GEBVs will be selected as parents.

### 2.2 Predicted cross value selection

The predicted cross value (PCV) calculates the probability that a pair of breeding parents will produce a gamete with desirable alleles at all specified loci by taking into account the recombination frequencies (18). This approach selects individuals based on their likelihood to produce an elite gamete by combining all desirable alleles. Since in a backcrossing scheme, individuals are always crossed with the recurrent parent, the PCV can be defined as the probability that each individual will produce an elite gamete.

Let *g* ∈ **B**^*L*×1^ denote a random gamete produced by a breeding parent. The PCV of an individual is calculated as follows:

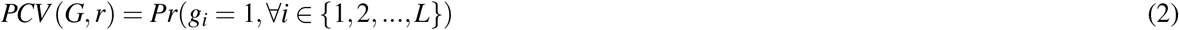

Here, *r* is the recombination frequency vector. To calculate this probability, the same water-pipe algorithm described in (14) is used. According to this approach, the positive individuals with highest PCVs will be selected as parents.

### 2.3 Look-ahead Monte Carlo selection

In this section we propose a novel probabilistic and heuristic driven search algorithm, look-ahead Monte Carlo (LMC) for parental selection. The underlying concept is to use Monte Carlo simulation for modeling uncertainty involved due to recombination events. Monte Carlo simulation is a technique that relies on repeated random sampling to obtain numerical results (27). This technique is often used in physical and mathematical problems and is most suited to be applied when it is impossible to obtain a closed-form expression or infeasible to apply a deterministic algorithm (28).

The look-ahead Monte Carlo algorithm for parental selection evaluates different selection decisions periodically during the learning phase by predicting the genetic distribution of the progeny of backcrosses after multiple generations using information of recombination events. This algorithm makes a trade-off between exploration and exploitation. It exploits the selection strategies that is found to be best until the current generation and also explores the alternative decisions to find out if they could replace the current best.

Figure 2 presents an overview of the LMC algorithm. For every individual in BCt population (e.g., BCt_*i*_), multiple random gametes are simulated according to the recombination frequencies. These gametes are narrowed down to the ones which have the desirable markers from the donor in the introgressed loci. Then, one of these positive gametes are selected randomly to form the next BC progeny (e.g., BC(t+1)_*i*_). This process is repeated until the target generation (BCT). Finally, individuals are evaluated based on their performance after selfing (BCTF1).

**Figure 2.**
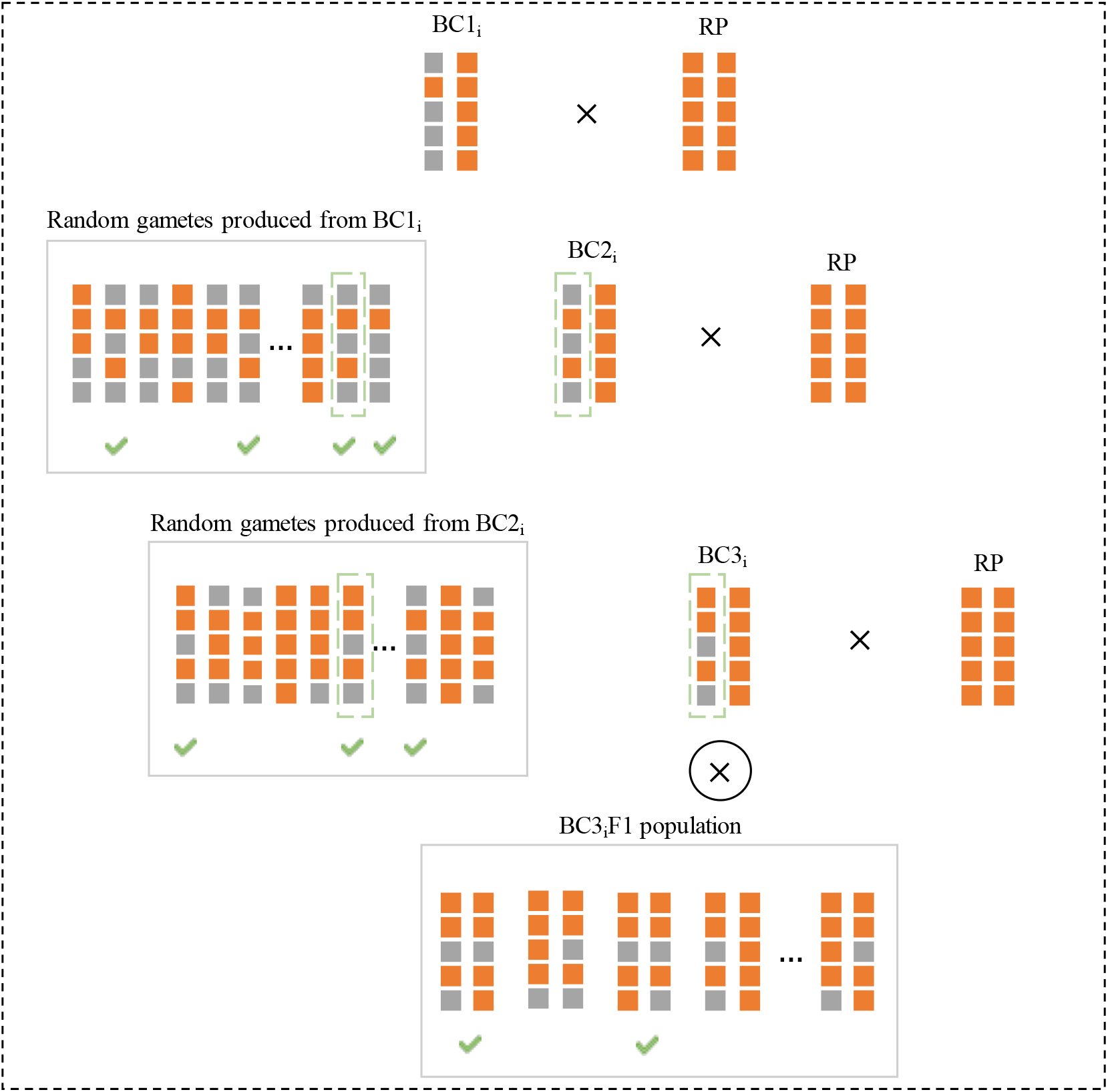
The Monte Carlo search for parental selection in trait introgression. For a deadline of T generations, we estimate the performance of BCTF1 individuals for a given selection strategy by searching across all possible paths.

In BCTF1, success can be defined as achieving certain amount of recovery percentage (e.g., *R* = 95) among positive individuals. Suppose the population size of the BCTF1 generation is *K* and n individuals with desirable markers have achieved at least R recovery percentage. Then 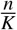 is the probability of getting a positive individual that has met the background recovery requirements. Since through backcross generations the gametes are selected randomly, this probability is estimating only one of the possible outcomes for individual *i* in BCt population. To have a reasonable approximation for the performance of progeny in BCTF1, the same process should be repeated multiple times. The objective of the LMC algorithm can then be calculated as:

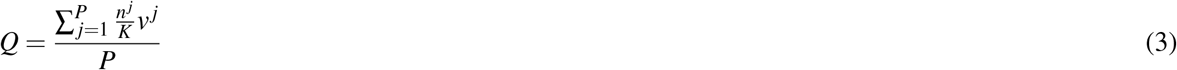

Where *v^j^* represents the maximum recovery percentage achieved in BCTF1 for the *j*^th^ round, and *P* is total rounds of repetition. According to LMC approach, individuals with highest *Q* values will be selected as the breeding parents.

## 3 Results

In this section, we first describe the data sets used in this case study, and then compare the proposed method with two existing selection methods in different scenarios of resources using computer simulation.

### 3.1 Data

Data contains donor and recipient’s genetic information and recombination frequencies. To explore the effect of having different initial genetic similarities between donor and recurrent parent, we considered three cases as demonstrated in Table 1. The genetic similarity is calculated based on the NEIs metric (29). Cases 1, 2, and 3 have low, moderate and high initial genetic similarities respectively.

**Table 1.**
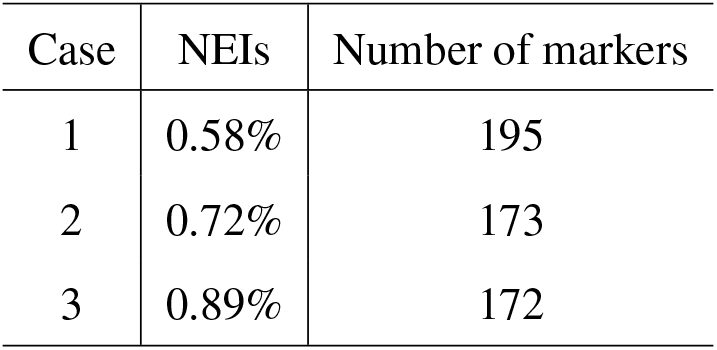
The description of data sets

The genetic information of donor and recurrent parent for these three cases are illustrated in Figure 3. For all cases, three markers should be integrated from the donor to the recurrent parent. Furtheremore, the recombination events are presented in Tables 3, 4, and 5 in the appendix.

**Figure 3.**
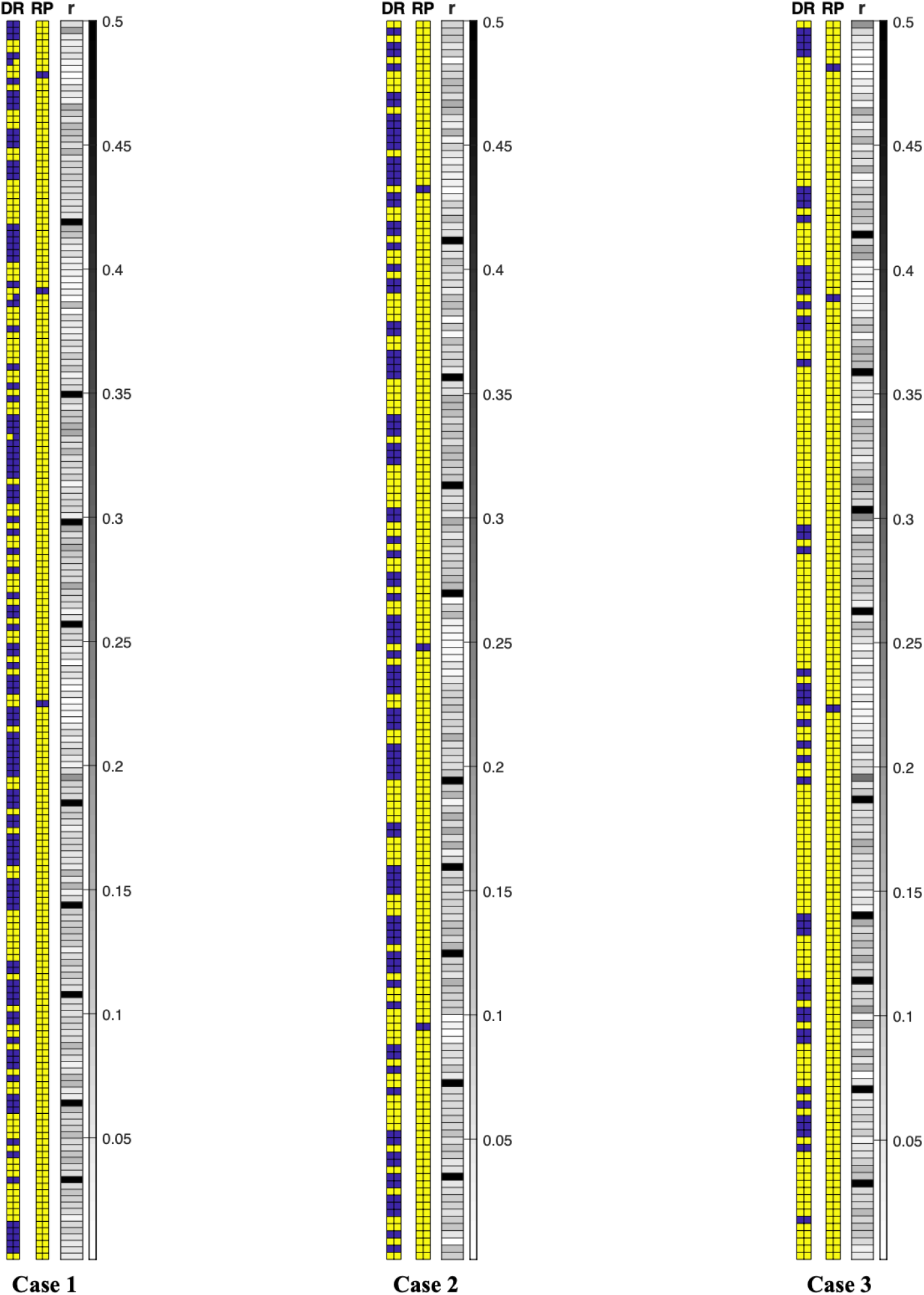
Donors and recurrent parents’ genetic information and recombination frequencies for three cases (DR: donor, RP: recurrent parent, r: recombination frequency). A yellow square is used to denote a favorable allele (“1”) and a purple square is used to denote an unfavorable allele (“0”). The gray charts are heat maps for recombination frequencies.

### 3.2 Simulation settings

In this study, simulations were conducted using MATLAB (R2019-a). One hundred independent simulation replicates were performed for each of the selection methods. Simulation has been performed for three generations of backcrosses followed by a selfing. The evaluation is based on the recovery percentage of individuals in BC3F1 generation. Figure 4 represents the flow chart for trait introgression pipeline.

**Figure 4.**
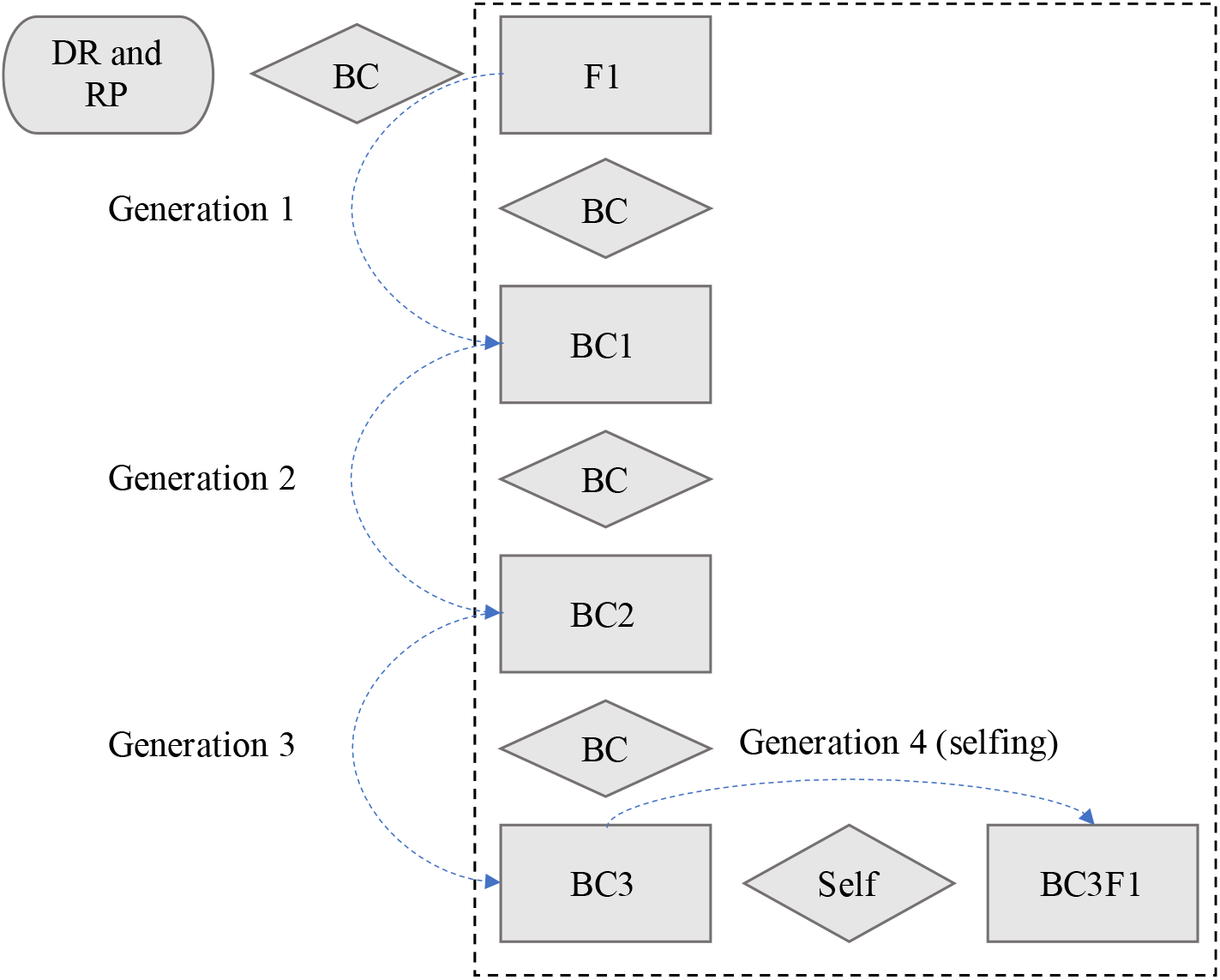
The trait introgression flowchart.

It is assumed that each cross makes 200 progeny and for each scenario the number of crosses remains the same through all generations (i.e. resources are distributed evenly among different generations).

### 3.3 Scenarios

To investigate the effect of resources on the performance among three methods, we have considered 2 different scenarios with varying numbers of crosses. In scenario 1, we are allocating limited resources by making 2 crosses in each generation whereas in scenario 2, we are allocating moderate resources by making 6 crosses in each generation (see Table 2). Scenario 1 more closely resembles what occurs in a commercial breeding program, namely, decision making with limited resources.

**Table 2.** Numbers of crosses in each generation for two different scenarios.

| Generation | Scenario 1<br>(limited resources) | Scenario 2<br>(moderate resources) |
| --- | --- | --- |
| BC1 | 2 | 6 |
| BC2 | 2 | 6 |
| BC3 | 2 | 6 |

### 3.4 Comparison of selection methods

Multiple trait introgression was studied using realistic maize data with three different selection methods, including GEBV, PCV, and LMC. We considered three cases with different genetic similarities and two different scenarios for resources.

Figure 5 presents the performance of three selection methods for one sample simulation. The histograms of background recovery percentage for positive individuals are demonstrated over BC1, BC2, BC3, and BC3F1 generations. All three methods start with the same BC1 population and then produce the next population based on different selection decisions. As expected, the background recovery improves from BC1 to BC3 for all selection methods. For this sample simulation, the (maximum, mean, minimum) recovery percentage in BC3 is (94, 90.61, 85), (94, 89.71, 84), (97, 94.07, 92) for GEBV, BPV, and LMC methods respectively which demonstrates improvement in recovery percentage when selection decisions are made using the LMC method.

**Figure 5.**
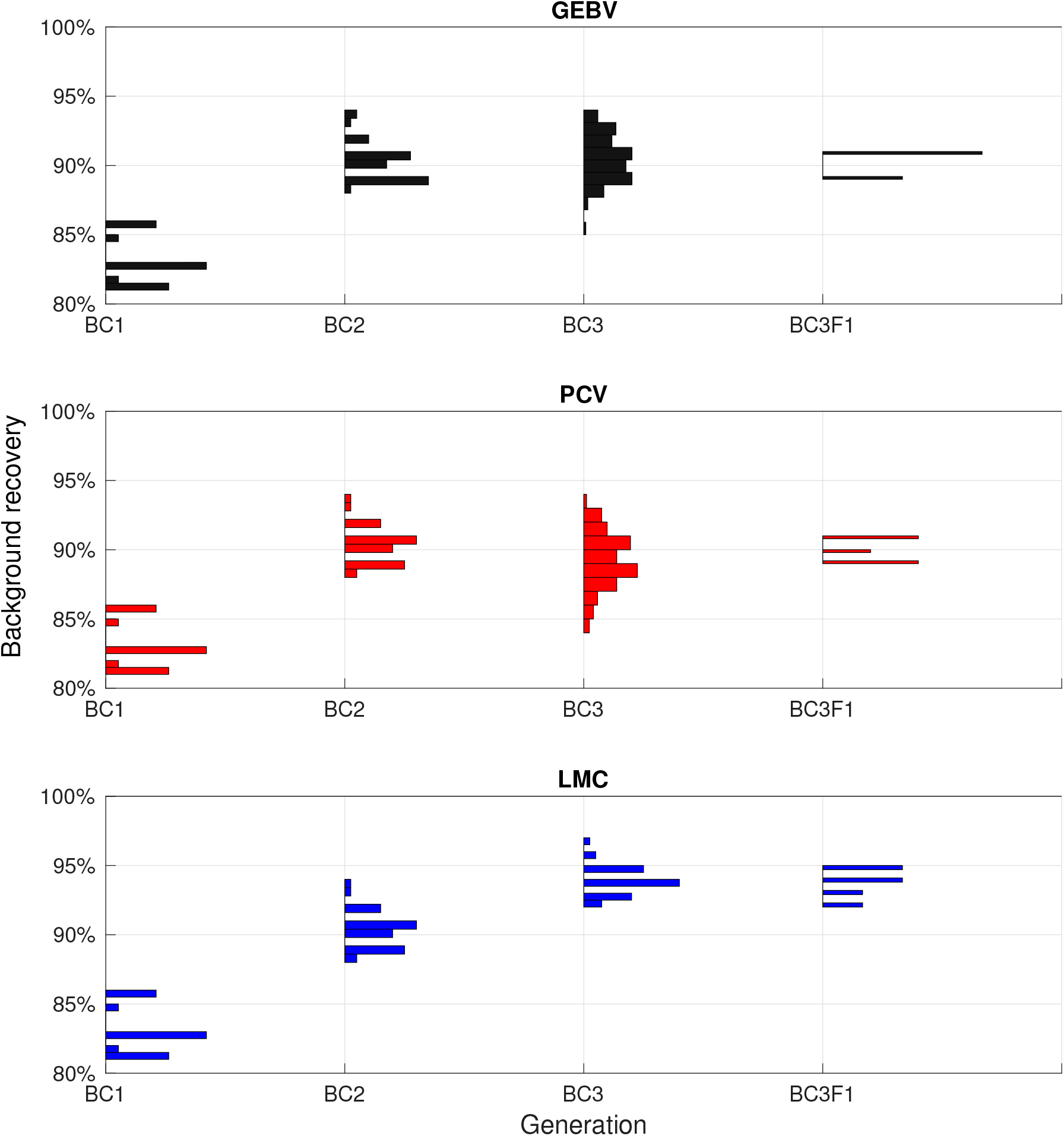
A sample simulation result for three different selection methods presenting histograms of population background recovery percentage over different generations. This simulation is performed for case 2, scenario 1.

It should be noted that the BC3F1 individuals should have all 6 alleles desirable in the three markers that are to be integrated from the donor (i.e. BC3F1 individuals are homozygous). However, the BC individuals are expected to have 3 desirable alleles total since their second chromosome is being inherited from the recurrent parent. This can explain why recovery percentage drops from BC3 to BC3F1. As demonstrated in Figure 5, for this sample simulation, the LMC method achieves 95% recovery in BC3F1, however the other two selection methods achieve maximum 91% recovery.

Figure 6 compares the cumulative distribution functions (CDFs) of maximum recovery percentage achieved in BC3 for three selection methods among 100 simulation replicates. The further toward the right direction of the figure a CDF curves, the better performance a method has. Take for example, point (97, 75) means that 75% of the simulations have achieved recovery percentage less than or equal to 97. In all cases and scenarios, the LMC method achieves higher recovery percentage.

**Figure 6.**
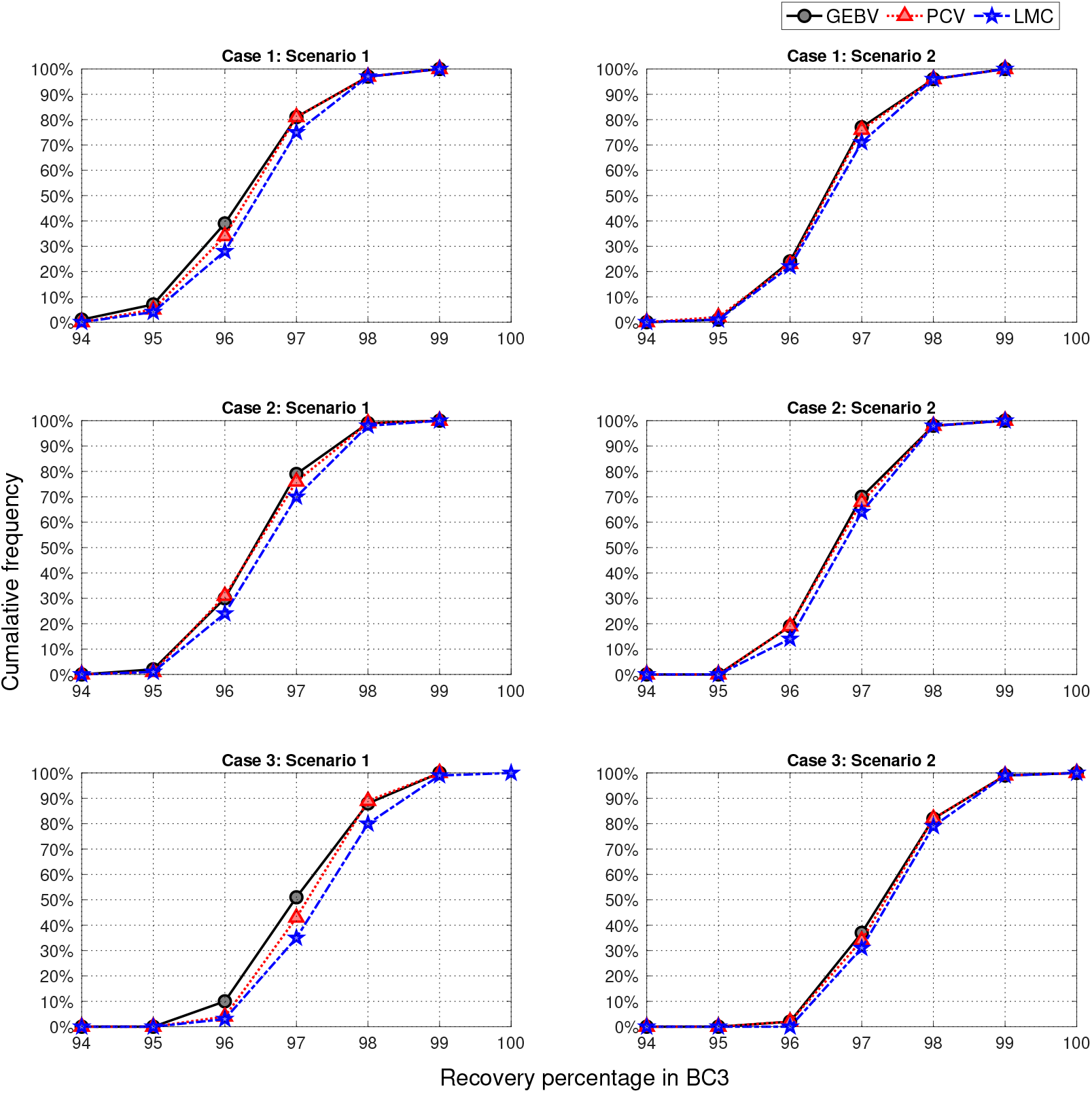
Cumulative distribution functions of population maximum in the BC3 for three cases and two different scenarios. Results are based on 100 simulation replicates.

For case 3 which has the highest genetic similarity between donor and recurrent parent, there was one simulation that resulted in having one individual in BC3 with all desirable traits (100% recovery percentage). Note that since this is a backcross generation, for this individual the second chromosome still lacks the desirable alleles from the donor. As expected, for each case, scenario 2 has better performance compared to scenario 1 since there are more resources available.

Figure 7 presents the box-plots of average recovery percentage of top 10 individuals in BC3 generation. For all cases and scenarios, the median value is higher when selection decisions are optimized using LMC method. Furthermore, PCV has generally higher median values than GEBV. The overall range of values is greater for LMC method (as shown by the distances between the ends of the two whiskers for each box-plot). The interquartile ranges are reasonably similar (as shown by the lengths of the boxes), except for case 2, scenario 1, where LMC has considerably higher range.

**Figure 7.**
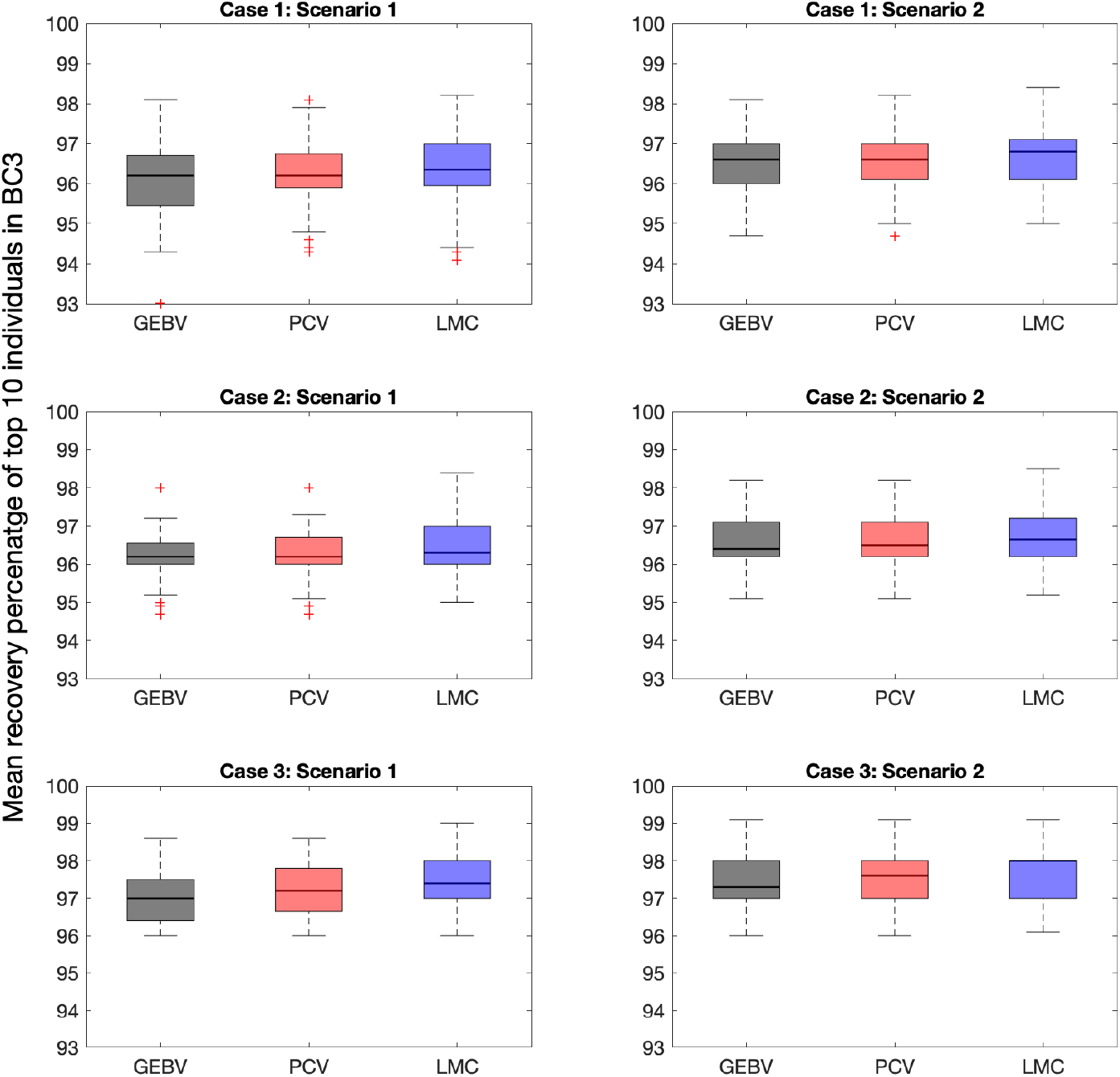
Box-plots of mean recovery percentage of top 10 individuals in BC3 for three selection methods. For each case and scenario, 100 simulation replicates are performed. The median values are demonstrated with a bold line.

Figure 8 compares the probability of success for three selection methods by evaluating recovery percentage of best individual in BC3F1. For example, point (0.8, 95) means that 80% of the simulations have achieved recovery percentage of 95 in the terminal generation. The curves with better performance are expected to be closer to the upper right corner of the plot.

**Figure 8.**
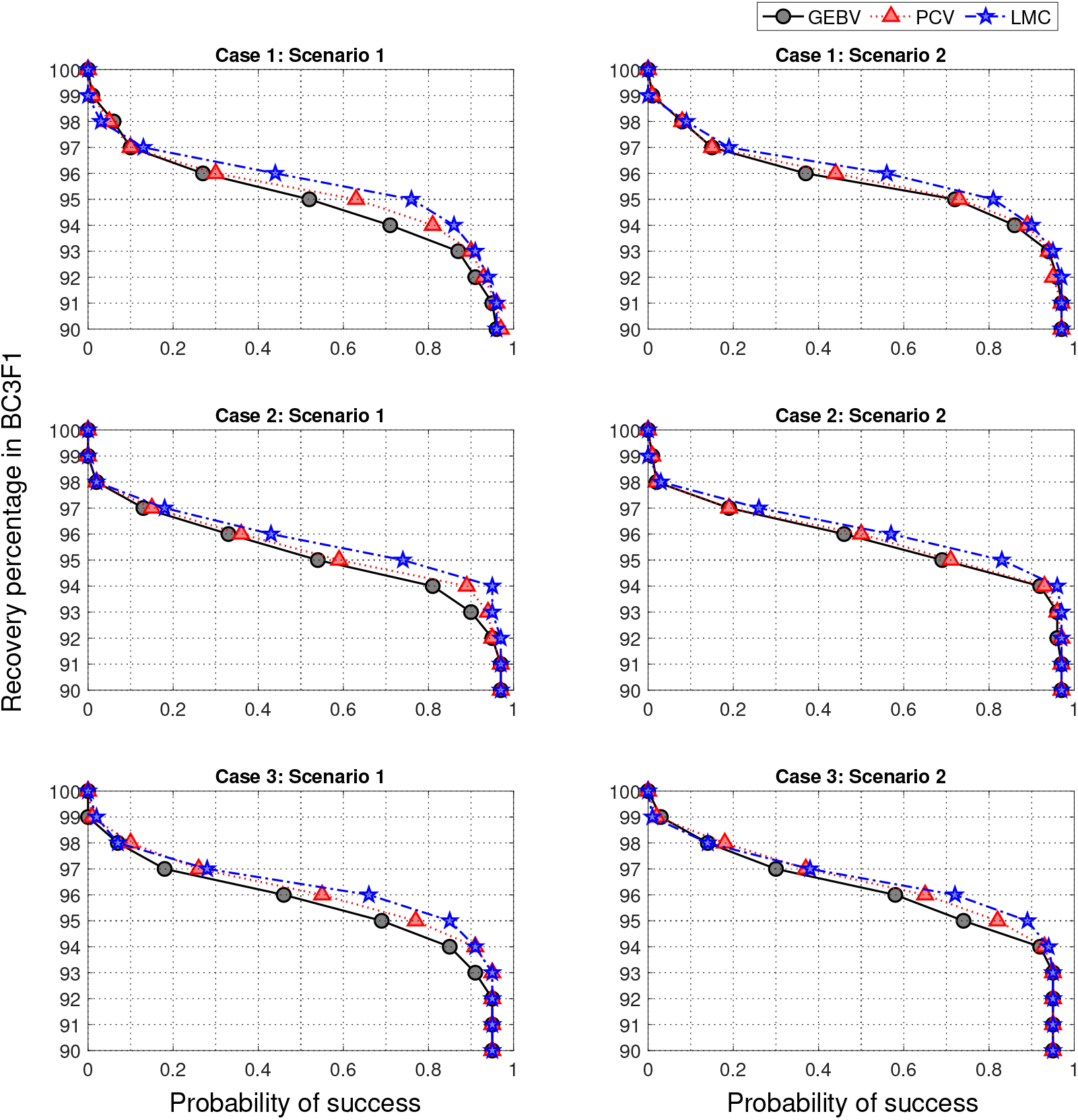
Probability of success in BC3F1 for three different selection approaches considering 3 cases of initial genetic similarity and 2 scenarios of resource allocation for 100 simulation replicates. The maximum recovery percentage of positive individuals in BC3F1 is first identified and then probability of success has been defined as the proportion of simulations that have achieved a certain recovery.

As expected, scenario 2 has generally higher probability of success compared to scenario 1 as more resources are used. Take for example, for case 2 the probability of achieving 95 percentage recovery with LMC method increases from 0.74 to 0.83 when having more resources. This probability also increases from 0.59 to 0.71 for PCV and from 0.54 to 0.69 for GEBV method. Furthermore, the probability of success increases from case 1 to 3 where there is more genetic similarity between the donor and parent. Take scenario 2 for example, the probabilities of having 95 percentage recovery when selection decisions are optimized using LMC method are 0.81, 0.83, 0.89 for cases 1, 2, and 3, respectively.

## 4 Discussion

Selection methods based on marker information make trait introgression more efficient and effective. When introgressing the desired traits form a donor to a recipient, background selection is the conventional selection approach that aims to recover the desired background genome. Recent advances in optimization and simulation techniques can help enhancing the efficiency of parental selection in breeding programs.

In this study, we introduced a new selection method, LMC, which has the potential to further improve the efficiency of breeding given limited time and resources by integrating operations research techniques and trait introgression. The proposed method was compared with existing selection methods in a simulation study using empirical maize data. Computational results demonstrate the improvements of the LMC method over two existing selection approaches, GEBV and PCV.

One of the advantages of the LMC method is being sensitive to the deadline. Unlike other selection methods that evaluate the performance based on only next generation, the LMC method relates the objective to the performance of individuals in the targeted generation. Another advantage of the LMC method is the trade-off between exploration and exploitation. When the look ahead process finds exploitation to contribute more to the final objective, the algorithm behaves in a greedy way to maximize performance. However, when the exploration is found to be more beneficial, the algorithm explores new possible outcomes.

The simulations in this study were designed based on practical considerations. The trait introgression pipeline included three backcross generations followed by a selfing so that selected individuals will be homozygous for the target trait. There is no absolute number for the number of backcrosses needed to be performed but generally between two to five backcrosses are performed in maize. The number of required generations can be determined based on the breeding objective and the resources invested at each generation (30). Intuitively, making more crosses and producing more progeny leads to a higher chance of creating desirable individuals, however the resources are limited and the breeding strategy should be customized based on available resources. Here, we considered two scenarios to represent both limited and moderate cases of resource availability. Scenario 1 limits the number of crosses to two in each generation where as scenario 2 allows six crosses. According to reproductive biology of maize, it is possible to obtain ≈ 200 seeds from a cross. Thus, we assumed each cross makes 200 progeny which means for scenarios 1 and 2, the population size of each generation becomes 400 and 1200 respectively. As expected, simulation results demonstrated that the probability of success increases when having more resources.

Future work should investigate optimizing the resource allocation strategies by spreading out the budget systematically among different generations. Also, this study investigated introgressing desirable alleles from a single donor, however desirable alleles can be carried by multiple donors. Hence, another direction that deserves investigation is to extend the LMC method for the cases with multiple donors. Moreover, we based our simulations on a single crop organism. Further simulations considering more diverse populations are necessary to demonstrate the general applicability of the proposed selection method.

## 5 Acknowledgements

This work was partially supported by Agriculture and Food Research Initiative Grant no.2017-67007-26175/Accession No. 1011702 from the USDA National Institute of Food and Agriculture. Additionally this work was partially supported by the National Science Foundation under the LEAP HI and GOALI programs (grant number 1830478). This work is also supported by the Plant Sciences Institute’s Faculty Scholars program at Iowa State University and Syngenta.

## 6 Appendix

Recombination frequencies for 3 data sets that were used in this study.

**Table 3.** Recombination frequencies for case 1. Each row refers to a marker and each column refers to a chromosome. The recombination frequencies for the three markers that should be integrated from the donor to the recipient are distinguished with an underline.

|  | Ch1 | Ch2 | Ch3 | Ch4 | Ch5 | Ch6 | Ch7 | Ch8 | Ch9 | Ch10 |
| --- | --- | --- | --- | --- | --- | --- | --- | --- | --- | --- |
| Marker1 | 0.0662 | 0.1441 | 0.0865 | 0.1588 | 0.0790 | 0.0915 | 0.0955 | 0.0662 | 0.1266 | 0.0722 |
| Marker2 | 0.1812 | 0.1122 | 0.0412 | 0.0705 | 0.0739 | 0.0955 | 0.0782 | 0.0636 | 0.0748 | 0.1176 |
| Marker3 | 0.0458 | 0.0574 | 0.1035 | 0.1067 | 0.0592 | 0.0627 | 0.1003 | 0.0610 | 0.0765 | 0.0915 |
| Marker4 | 0.0357 | 0.0394 | 0.0979 | 0.1615 | 0.0282 | 0.0253 | 0.1152 | 0.0874 | 0.0906 | 0.0857 |
| Marker5 | 0.0512 | 0.0671 | 0.1533 | 0.1051 | 0.0679 | 0.0739 | 0.0955 | 0.0857 | 0.1325 | 0.0731 |
| Marker6 | 0.0627 | 0.0310 | 0.1682 | 0.0849 | 0.0060 | 0.0857 | 0.0987 | 0.0592 | 0.0773 | 0.0234 |
| Marker7 | 0.0167 | 0.0347 | 0.0697 | 0.0748 | 0.0610 | 0.0857 | 0.0263 | 0.0636 | 0.0636 | 0.0739 |
| Marker8 | <u>0.0089</u> | 0.0282 | 0.0697 | 0.0748 | 0.0557 | 0.0503 | 0.0688 | 0.1398 | 0.1145 | 0.1003 |
| Marker9 | 0.0050 | 0.0158 | 0.1413 | 0.0636 | 0.0215 | 0.0565 | 0.1176 | 0.0840 | 0.1347 | 0.0592 |
| Marker10 | 0.0089 | <u>0.0089</u> | 0.0565 | 0.1688 | 0.0070 | 0.0412 | 0.0653 | 0.0557 | 0.0748 | 0.0412 |
| Marker11 | 0.0662 | 0.0253 | 0.0857 | 0.0815 | 0.0530 | 0.1377 | 0.1090 | 0.0282 | 0.0653 | 0.0494 |
| Marker12 | 0.0476 | 0.0010 | 0.0583 | 0.0756 | <u>0.0020</u> | 0.0412 | 0.0679 | 0.0512 | 0.0476 |  |
| Marker13 | 0.0244 | 0.1333 | 0.0939 | 0.0739 | 0.0196 | 0.1484 | 0.0627 | 0.1281 | — | — |
| Marker14 | 0.1581 | 0.0040 | 0.0244 | 0.0244 | 0.0119 | 0.0301 | — | 0.1377 | — | — |
| Marker15 | 0.1051 | 0.0722 | 0.0618 | 0.0512 | 0.0010 | 0.0645 | — | 0.0440 | — | — |
| Marker16 | 0.0494 | 0.0430 | 0.0636 | — | 0.0375 | — | — | 0.0394 | — | — |
| Marker17 | 0.1391 | 0.0282 | 0.0832 | — | 0.0731 | — | — | — | — | — |
| Marker18 | 0.1191 | 0.0476 | 0.0748 | — | 0.0301 | — | — | — | — | — |
| Marker19 | 0.0995 | 0.0782 | 0.0366 | — | 0.0583 | — | — | — | — | — |
| Marker20 | 0.0662 | 0.0476 | — | — | 0.0931 | — | — | — | — | — |
| Marker21 | 0.1470 | 0.0947 | — | — | 0.0421 | — | — | — | — | — |
| Marker22 | 0.0539 | 0.0618 | — | — | 0.0319 | — | — | — | — | — |
| Marker23 | 0.0931 | 0.1075 | — | — | 0.0653 | — | — | — | — | — |
| Marker24 | 0.0765 | 0.0253 | — | — | 0.2045 | — | — | — | — | — |
| Marker25 | 0.1011 | 0.0583 | — | — | 0.0874 | — | — | — | — | — |
| Marker26 | 0.0748 | 0.0722 | — | — | 0.0782 | — | — | — | — | — |
| Marker27 | 0.0714 | — | — | — | 0.0739 | — | — | — | — | — |
| Marker28 | 0.0782 | — | — | — | — | — | — | — | — | — |
| Marker29 | 0.0565 | — | — | — | — | — | — | — | — | — |
| Marker30 | 0.0592 | — | — | — | — | — | — | — | — | — |
| Marker31 | 0.0548 | — | — | — | — | — | — | — | — | — |

**Table 4.** Recombination frequencies for case 2.

|  | Ch1 | Ch2 | Ch3 | Ch4 | Ch5 | Ch6 | Ch7 | Ch8 | Ch9 | Ch10 |
| --- | --- | --- | --- | --- | --- | --- | --- | --- | --- | --- |
| Marker1 | 0.0799 | 0.0485 | 0.0674 | 0.0733 | 0.0028 | 0.0910 | 0.0757 | 0.0855 | 0.0770 | 0.0524 |
| Marker2 | 0.0953 | 0.0626 | 0.0618 | 0.0614 | 0.0801 | 0.1472 | 0.1118 | 0.0966 | 0.0724 | 0.0671 |
| Marker3 | 0.1057 | 0.1189 | 0.1467 | 0.0990 | 0.1353 | 0.0246 | 0.0587 | 0.0498 | 0.0793 | 0.0959 |
| Marker4 | 0.0484 | 0.0868 | 0.1368 | 0.0406 | 0.0279 | 0.0814 | 0.0870 | 0.1510 | 0.0721 | 0.0914 |
| Marker5 | 0.0911 | 0.0659 | 0.1306 | 0.1580 | 0.0236 | 0.1506 | 0.0928 | 0.1049 | 0.0894 | 0.0567 |
| Marker6 | 0.0040 | 0.0693 | 0.0752 | 0.1174 | 0.0130 | 0.0737 | 0.0861 | 0.0888 | 0.0820 | 0.1009 |
| Marker7 | 0.0792 | 0.0392 | 0.1079 | 0.0551 | 0.0113 | 0.1709 | 0.0640 | 0.0915 | 0.1219 | 0.1118 |
| Marker8 | 0.0815 | 0.1110 | 0.1199 | 0.1113 | **0.0150** | 0.0663 | 0.1220 | 0.0966 | 0.0598 | 0.0510 |
| Marker9 | 0.1409 | 0.1297 | 0.0646 | 0.0625 | 0.0129 | 0.1496 | 0.1310 | 0.0016 | 0.1178 | 0.0025 |
| Marker10 | 0.1164 | 0.0997 | 0.1395 | 0.0790 | 0.0484 | 0.0285 | 0.0739 | **0.0258** | 0.0979 | 0.1216 |
| Marker11 | 0.0670 | 0.0086 | 0.1265 | 0.0975 | 0.0115 | 0.0650 | 0.1451 | 0.0033 | 0.0545 | 0.0966 |
| Marker12 | 0.1578 | 0.1076 | 0.1031 | 0.0975 | 0.0481 | — | — | 0.0146 | 0.0595 | — |
| Marker13 | 0.1389 | 0.0115 | 0.0823 | 0.1265 | 0.0775 | — | — | 0.0645 | — | — |
| Marker14 | 0.0542 | 0.1247 | 0.1043 | 0.1357 | 0.0922 | — | — | 0.0653 | — | — |
| Marker15 | 0.0356 | 0.0729 | — | — | 0.0899 | — | — | 0.0541 | — | — |
| Marker16 | 0.1474 | 0.0843 | — | — | 0.1107 | — | — | 0.1075 | — | — |
| Marker17 | 0.0213 | 0.0833 | — | — | 0.0716 | — | — | 0.0450 | — | — |
| Marker18 | 0.0938 | 0.0365 | — | — | 0.0923 | — | — | — | — | — |
| Marker19 | 0.0527 | — | — | — | 0.0657 | — | — | — | — | — |
| Marker20 | 0.0311 | — | — | — | 0.1572 | — | — | — | — | — |
| Marker21 | 0.0490 | — | — | — | 0.0714 | — | — | — | — | — |
| Marker22 | 0.0269 | — | — | — | 0.0853 | — | — | — | — | — |
| Marker23 | **0.0279** | — | — | — | 0.0670 | — | — | — | — | — |
| Marker24 | 0.0080 | — | — | — | 0.0653 | — | — | — | — | — |
| Marker25 | 0.0257 | — | — | — | 0.1101 | — | — | — | — | — |
| Marker26 | 0.0890 | — | — | — | — | — | — | — | — | — |
| Marker27 | 0.0490 | — | — | — | — | — | — | — | — | — |
| Marker28 | 0.1303 | — | — | — | — | — | — | — | — | — |
| Marker29 | 0.0515 | — | — | — | — | — | — | — | — | — |
| Marker30 | 0.0636 | — | — | — | — | — | — | — | — | — |

**Table 5.** Recombination frequencies for case 3.

|  | Ch1 | Ch2 | Ch3 | Ch4 | Ch5 | Ch6 | Ch7 | Ch8 | Ch9 | Ch10 |
| --- | --- | --- | --- | --- | --- | --- | --- | --- | --- | --- |
| Marker1 | 0.2098 | 0.0722 | 0.0653 | 0.2332 | 0.0421 | 0.0697 | 0.1635 | 0.0697 | 0.0338 | 0.0756 |
| Marker2 | 0.0739 | 0.1708 | 0.0384 | 0.0756 | 0.1333 | 0.0592 | 0.0756 | 0.0722 | 0.0557 | 0.0748 |
| Marker3 | 0.0476 | 0.1695 | 0.0790 | 0.0503 | 0.0890 | 0.0906 | 0.1035 | 0.1027 | 0.0653 | 0.0874 |
| Marker4 | 0.0440 | 0.0128 | 0.0530 | 0.1615 | 0.0592 | 0.0583 | 0.1377 | 0.1888 | 0.1019 | 0.1355 |
| Marker5 | 0.0030 | 0.0225 | 0.0512 | 0.1003 | 0.0119 | 0.0739 | 0.0882 | 0.0234 | 0.0539 | 0.0865 |
| Marker6 | **<u>0.0060</u>** | 0.0282 | 0.0060 | 0.1214 | 0.0574 | 0.1648 | 0.1662 | 0.1760 | 0.0987 | 0.1122 |
| Marker7 | 0.0050 | 0.0158 | 0.1512 | 0.0731 | 0.0030 | 0.1251 | 0.0679 | 0.0565 | 0.0244 | 0.1051 |
| Marker8 | 0.0089 | **<u>0.0089</u>** | 0.1152 | 0.1129 | 0.0592 | 0.0583 | 0.1027 | 0.0467 | 0.0557 | 0.0756 |
| Marker9 | 0.0347 | 0.0263 | 0.1075 | 0.0955 | 0.0375 | 0.0898 | — | 0.0177 | 0.0601 | 0.0310 |
| Marker10 | 0.0329 | 0.0291 | 0.0756 | 0.1183 | 0.0449 | 0.0688 | — | 0.1533 | 0.0898 | 0.0476 |
| Marker11 | 0.0874 | 0.0206 | 0.0688 | 0.1043 | 0.0070 | 0.0548 | — | 0.0548 | 0.1405 | — |
| Marker12 | 0.0196 | 0.0963 | 0.0291 | 0.0679 | 0.0530 | 0.0799 | — | 0.1413 | 0.1137 | — |
| Marker13 | 0.1601 | 0.1244 | 0.0923 | 0.0329 | **<u>0.0020</u>** | 0.0301 | — | 0.0020 | — | — |
| Marker14 | 0.0857 | 0.0099 | 0.1051 | — | 0.0196 | 0.0601 | — | 0.0485 | — | — |
| Marker15 | 0.0244 | 0.1145 | 0.1831 | — | 0.0128 | 0.0244 | — | — | — | — |
| Marker16 | 0.0824 | 0.1628 | 0.1427 | — | 0.0384 | — | — | — | — | — |
| Marker17 | 0.1498 | 0.1296 | 0.0832 | — | 0.0440 | — | — | — | — | — |
| Marker18 | 0.0375 | 0.1289 | 0.0748 | — | 0.0310 | — | — | — | — | — |
| Marker19 | 0.0815 | — | — | — | 0.0824 | — | — | — | — | — |
| Marker20 | 0.0618 | — | — | — | 0.0263 | — | — | — | — | — |
| Marker21 | 0.1574 | — | — | — | 0.0739 | — | — | — | — | — |
| Marker22 | 0.0119 | — | — | — | 0.0705 | — | — | — | — | — |
| Marker23 | 0.0530 | — | — | — | 0.2767 | — | — | — | — | — |
| Marker24 | 0.1114 | — | — | — | 0.0512 | — | — | — | — | — |
| Marker25 | 0.0773 | — | — | — | 0.1051 | — | — | — | — | — |
| Marker26 | 0.1168 | — | — | — | — | — | — | — | — | — |
| Marker27 | 0.1281 | — | — | — | — | — | — | — | — | — |
| Marker28 | 0.1137 | — | — | — | — | — | — | — | — | — |
| Marker29 | 0.0539 | — | — | — | — | — | — | — | — | — |

